# Preparatory delta phase response is correlated with naturalistic speech comprehension performance

**DOI:** 10.1101/827584

**Authors:** Jiawei Li, Bo Hong, Guido Nolte, Andreas K. Engel, Dan Zhang

## Abstract

While human speech comprehension is thought to be an active process that involves top-down predictions, it remains unclear how predictive information is used to prepare for the processing of upcoming speech information. We aimed to identify the neural signatures of the preparatory processing of upcoming speech. Participants selectively attended to one of two competing naturalistic, narrative speech streams, and a temporal response function (TRF) method was applied to derive event-related-like neural responses from electroencephalographic data. The phase responses to the attended speech at the delta band (1–4 Hz) were correlated with the comprehension performance of individual participants, with a latency of -200–0 ms before onset over the fronto-central and left-lateralized parietal regions. The phase responses to the attended speech at the alpha band also correlated with comprehension performance, but with a latency of 650–980 ms post-onset over fronto-central regions. Distinct neural signatures were found for the attentional modulation, taking the form of TRF-based amplitude responses at a latency of 240–320 ms post-onset over the left-lateralized fronto-central and occipital regions. Our findings reveal how the brain gets prepared to process an upcoming speech in a continuous, naturalistic speech context.

## 1. Introduction

Humans can effectively comprehend complex and rapidly changing speech in challenging conditions, e.g., in a cocktail party scenario with multiple competing speech streams and high background noise. To achieve such a capacity, the human brain is equipped with an efficient neural architecture that is dedicated to bottom-up processing of perceived speech information, from the low-level acoustics, to the phoneme, syllable, and sentence levels (Pisoni and Luce 1987; DeWitt and Rauschecker 2012; Friederici 2012; Hickok 2012; Verhulst et al. 2018). In recent years, increasing evidence has also suggested that human speech comprehension is an active process that involves top-down predictions (Rao and Ballard 1999; Federmeier 2007; Arnal et al. 2011; Hickok et al. 2011; Kutas and Federmeier 2011; Fries 2015; Tian et al. 2018). In the cocktail party scenario, it is believed that a listener should continuously predict what their attended speaker is going to say next in order to efficiently understand the corresponding speech (Cherry 1953; Ding and Simon 2012; Zion Golumbic, Cogan, et al. 2013; O’Sullivan et al. 2015; Bednar and Lalor 2020).

Although the idea of prediction in human speech comprehension is gaining popularity, it remains unclear how the brain gets prepared for the processing of upcoming speech information. The available findings on prediction in speech, however, are not sufficient to determine the neural mechanisms underlying preparation. For instance, the classic studies of active speech prediction have mainly focused on neural activity in response to prediction errors. Event-related potential (ERP) components such as the N400 and P600 are frequently reported when the perceived word violates semantic and syntactic congruency of the preceding speech context, respectively (Lau et al. 2008; Kutas and Federmeier 2011; Van Petten and Luka 2012; Wang et al. 2018). These ERP components normally occur >400 ms after the presentation of the perceived speech, and so provide only indirect support for the preparatory process.

Recent studies have reported evidence of the brain’s pre-activation before the onset of the upcoming speech. Some studies have focused on preparatory attentional orientation to specific acoustic features (e.g., spatial location, pitch, etc.) in speech or general auditory tasks (Hill and Miller 2010; Lee et al. 2013; Holmes et al. 2016, 2018; ElShafei et al. 2018; Nolden et al. 2019). It has been demonstrated that an external cue (e.g., a visual symbol) prior to the onset of the auditory or speech stimuli could elicit distinct cue-specific neural responses, depending on the cue-instructed attentional task. In the meanwhile, researchers have reported pre-activations that are more specific for speech processing (DeLong et al. 2005; Dikker and Pylkkänen 2013; Söderström et al. 2016, 2018): event-related neural responses to a preceding speech unit (e.g., words) were found to be informative about possible upcoming speech units in the continuous speech materials (e.g., sentences). While these speech-related pre-activations could reflect the brain’s preparation for processing the upcoming speech, they were represented by event-related responses to either attention-related cues or preceding speech units. Therefore, the possible findings are dependent upon the nature of these preceding events, which are substantially different from the upcoming, to-be-processed speech unit. More importantly, it could be further argued that these pre-activations might only reflect a neural processing phase prior to the actual preparatory speech processing, and there should be a more direct preparatory phase during the processing of the upcoming speech. Ideally, the most direct preparatory processing should be linked to the to-be-processed speech but occur before its onset.

While this direct evidence has not been investigated in the speech domain, studies on general sensory processing have provided support for the existence of such a preparatory process. In the visual domain, the amplitude and phase of pre-stimulus oscillatory activities, especially in the alpha band, have been reported to have a significant impact on subsequent perceptual consequences, (Van Dijk et al. 2008; Kok et al. 2017; Harris et al. 2018; Galindo-Leon et al. 2019; Rassi et al. 2019), such as threshold-level perception, attention orientation, visual search performance, etc. Auditory information processing has also been shown to be affected by pre-stimulus oscillations, mainly at lower frequency bands such as theta and delta (Ng et al. 2012; Kayser et al. 2016; Zoefel et al. 2018). However, these studies have mainly employed simple and abstract sensory stimuli, therefore limited in explaining possible neural mechanisms underlying the preparatory processing of human speech. Recently, there is an emergence of naturalistic stimuli in auditory studies (Sonkusare et al. 2019). Researchers adopted naturalistic, continuous stimuli (e.g., poetry, long sentences) as the stimulus (Etard and Reichenbach 2019; Donhauser and Baillet 2020; Teng et al. 2020). Naturalistic speech presents listeners with a variety and multitude of different linguistic contents (Alexandrou et al. 2020), catering for different levels of preparation. Thus, our study utilized naturalistic and continuous materials to study the preparatory processing.

One crucial issue that needs to be considered is the possible dependence of the preparatory process on top-down selective attention. As attention regulates the processing of the input sensory information, it can be expected to affect prediction and consequently preparation (Schröger, Kotz, et al. 2015; Schröger, Marzecová, et al. 2015). A number of recent studies have indeed reported attention-dependent neural responses to prediction errors (Kok et al. 2012; Auksztulewicz and Friston 2015; Hisagi et al. 2015; Marzecová et al. 2017; Smout et al. 2019), with ongoing debates on the direction of the interplay and the involved sensory processing stages. Most of these studies have been conducted within the visual domain, with limited exploration in the auditory domain, let alone speech processing.

The present study aimed to identify neural signatures that directly reflect the preparatory processing of human speech. Naturalistic, narrative speech materials were presented auditorily to the participants with a 60-channel electroencephalogram (EEG) recording; this procedure is believed to be of high ecological validity and thus to provide necessary contextual information for the engagement of top-down prediction and therefore preparation (Rao and Ballard 1999; Friston 2005; Federmeier 2007; Jehee and Ballard 2009). A cocktail party paradigm was used to introduce a complex perceptual environment that imposed further demands on prediction and preparation (Cherry 1953; Ding and Simon 2012; Zion Golumbic, Cogan, et al. 2013; O’Sullivan et al. 2015; Broderick et al. 2018). To characterize the neural responses to the continuous, naturalistic speech streams, a temporal response function (TRF) method was used to derive event-related-like neural responses from the EEG signals, based on the speech power envelop of both the attended and the unattended speech streams (Lalor et al. 2006; Crosse et al. 2016). The TRF-based responses revealed the temporal dynamics of neural activities underlying human speech processing, and the responses with latencies earlier than the onset of speech power envelop fluctuations are regarded to represent the preparatory phase. Specifically, the TRF-based responses were further decomposed into amplitude and phase responses, as amplitude and phase have been proposed to play unique roles in networks underlying human cognition (Bonnefond and Jensen 2012; Engel et al. 2013; Fries 2015). Following the studies on the perceptual influence of pre-stimulus neural activities (Smith et al. 2006; Iemi et al. 2019; Rassi et al. 2019; Avramiea et al. 2020), we were interested in whether the TRF-based responses at the preparatory stage could be correlated to speech comprehension performance, as measured by speech-content-related questionnaires, and how amplitude and phase responses contributed to speech preparation. With the employment of the cocktail party paradigm, we also addressed the issue about the attention-dependency of the to-be-explored performance-related preparatory activities. Our study is expected to reveal the neural mechanisms underlying how the brain gets prepared to process an upcoming speech in a continuous, naturalistic speech context.

## 2. Materials and Methods

### 2.1. Ethics statement

The study was conducted in accordance with the Declaration of Helsinki and was approved by the local Ethics Committee of Tsinghua University. Written informed consent was obtained from all participants.

### 2.2. Experimental model and participant details

Twenty college students (10 females; mean age: 24.7 years; range: 20–43 years) from Tsinghua University participated in the study as paid volunteers. The sample size (N = 20) was decided following previous TRF-based studies on human speech processing (Di Liberto et al. 2015; Mirkovic et al. 2015; Broderick et al. 2018). All participants were native Chinese speakers, reported having normal hearing, and normal or corrected-to-normal vision.

### 2.3. Stimuli

The speech stimuli were recorded from two male speakers using the microphone of an iPad2 mini (Apple Inc., Cupertino, CA) at a sampling rate of 44,100 Hz. The speakers were college students from Tsinghua University, who had more than four years of professional training in broadcasting. Both speakers were required to tell 28 1-min narrative stories in Mandarin Chinese; the stories were either those about daily-life topics recommended by the experimenter and told by the speaker improvising on their own (14 stories), or those selected from the National Mandarin Proficiency Test (14 stories). The speakers were presented with the recommended topic or story materials on the computer screen. They were allowed to prepare for as long as required before telling the story (usually ∼3 min). When they were ready, the speakers pressed the SPACE key on the computer keyboard and the recording began with the presentation of three consecutive pure-tone beep sounds at 1000 Hz (duration: 1000 ms; inter-beep interval: 1500 ms). The beep sounds served as the event marker to synchronize the speech audios in the main experiment, in which two speech streams were presented simultaneously. The speakers were asked to start speaking as soon as the third beep had ended (within around 3 sec). The speakers were allowed to start the recording again if the audio did not meet the requirements of either the experimenter or the speakers themselves (which mainly concerned speech coherence). The actual speaking time per story ranged from 51 to 76 sec.

Two four-choice questions per story were then prepared by the experimenter and two college students who were familiar with comprehension performance assessment. These questions and the corresponding choices concerned story details that required significant attentional efforts. For instance, one question following a story about one’s hometown was, “What is the most dissatisfying thing about the speaker’s hometown? (推测讲述人对于家乡最不满意的地方在于?)”, and the four choices were A) There is no heating in winter; B) There are no hot springs in summer; C) There is no fruit in autumn; D) There are no flowers in spring (A. 冬天没暖气;B. 夏天没温泉;C. 秋天没水果;D. 春 天 没 鲜 花). Both the speech audio and corresponding questions are available for downloads.

### 2.4. Experimental procedure

The main experiment consisted of 4 blocks of 7 trials per block. In each trial, two narrative stories were presented simultaneously, one to the left ear and the other to the right ear. The participants were instructed to attend to one spatial side. The two speech streams within each trial were from the two different male speakers to facilitate selective attention. Considering the possible duration difference between the two audio streams, the trial ended after the longer speech audio had ended. Each trial began when participants pressed the SPACE key on the computer keyboard. Participants were instructed which side to attend to by plain text (“Please pay attention to the [LEFT/RIGHT]”) displayed on the computer screen. A white fixation cross was also displayed throughout the trial. The speech stimuli were played immediately after the keypress, and were preceded by the three beep sounds to allow participants to prepare. At the end of each trial, four questions (two for each story) were presented sequentially in a random order on the computer screen, and the participants made their choices using the computer keyboard. After completing these questions, participants scored their attention level of the attended stream, the experienced difficulty of performing the attention task, and the familiarity with the attended material using three 10-point Likert scales. No feedback was given to the participants about their performance during the experiment. Throughout the trial, participants were required to maintain visual fixation on the fixation cross while listening to the speech and to minimize eye blinks and all other motor activity. The participants were recommended to take a short break (of around 1 min) after every trial within one block, and a long break (no longer than 10 min) between blocks.

The to-be-attended side was fixed within each block (two blocks for attending to the left side and two for attending to the right side). Within each block, the speaker identity remained unchanged for the left and right sides. In this way, the to-be-attended spatial side and the corresponding speaker identity were balanced within the participant, with seven trials per side for both speakers. The assignment of the stories to the four blocks was randomized across the participants.

The experiment was carried out in a sound-attenuated, dimly lit, and electrically shielded room. The participants were seated in a comfortable chair in front of a 19.7-inch Lenovo LT2013s Wide LCD monitor. The viewing distance was approximately 60 cm. The experimental procedure was programmed in MATLAB using the Psychophysics Toolbox 3.0 extensions (Brainard and Brainard 1997). The speech stimuli were delivered binaurally via an air-tube earphone (Etymotic ER2, Etymotic Research, Elk Grove Village, IL, USA) to avoid possible electromagnetic interferences from auditory devices. The volume of the audio stimuli was adjusted to be at a comfortable level that was well above the auditory threshold. Furthermore, the speech stimuli driving the earphone were used as an analog input to the EEG amplifier through one of its bipolar inputs together with the EEG recordings. In this way, the audio and the EEG recordings were precisely synchronized, with a maximal delay of 1ms (at a sampling rate of 1000 Hz).

### 2.5. Data acquisition and pre-processing

EEG was recorded from 60 electrodes (FP1/2, FPZ, AF3/4, F7/8, F5/6, F3/4, F1/2, FZ, FT7/8, FC5/6, FC3/4, FC1/2, FCZ, T7/8, C5/6, C3/4, C1/2, CZ, TP7/8, CP5/6, CP3/4, CP1/2, CPZ, P7/8, P5/6, P3/4, P1/2, PZ, PO7/8, PO5/6, PO3/4, POZ, Oz, and O1/2), which were referenced to an electrode between Cz and CPz, with a forehead ground at Fz. A NeuroScan amplifier (SynAmp II, NeuroScan, Compumedics, USA) was used to record EEG at a sampling rate of 1000 Hz. Electrode impedances were kept below 10 kOhm for all electrodes.

The recorded EEG data were first notch filtered to remove the 50 Hz powerline noise and then subjected to an artifact rejection procedure using independent component analysis. Independent components (ICs) with large weights over the frontal or temporal areas, together with a corresponding temporal course showing eye movement or muscle movement activities, were removed. The remaining ICs were then back-projected onto the scalp EEG channels, reconstructing the artifact-free EEG signals. While the relatively long duration of the speech trials in the present study (∼1 minute per story, see Experimental procedure) has made it more difficult for the participants to avoid inducing movement-related artifacts as compared to the classical ERP-based studies, a temporally continuous, non-interrupted EEG segment per trial was preferred for the employment of the TRF method. Therefore, any ICs with artifact-like EEG activities for more than 20% of the trial time (i.e. ∼12 sec) were rejected, leading to around 4–11 ICs rejected per participant. The cleaned EEG data was used for the TRF analysis without any further artifact rejection procedures. Then the EEG signals were re-referenced to a common average reference.

Next, the EEG data were segmented into 28 trials according to the markers representing speech onsets. The analysis window for each trial extended from 10 to 55 s (duration: 45 s) to avoid the onset and the offset of the stories.

### 2.6. Temporal response function modeling

The analysis workflow is shown in Figure 1. The neural responses to the speech stimuli were characterized using a temporal response function (TRF)-based modeling method. The TRF response describes the impulse response to fluctuations of an input signal, and is based on system identification theories (Lalor et al. 2006; Crosse et al. 2016). We used the power envelope of the speech signal as the input signal required by TRF, which has been demonstrated to be a valid index by which to extract speech-related neural responses (Ding and Simon 2012; Mirkovic et al. 2015; O’Sullivan et al. 2015; Bednar and Lalor 2018; Broderick et al. 2018; Huang et al. 2018). TRF-based responses before the onset of the speech power envelope fluctuations are considered to represent preparatory activity, whereas post-onset responses reflect post-processing of the speech stream.

**Figure 1.**
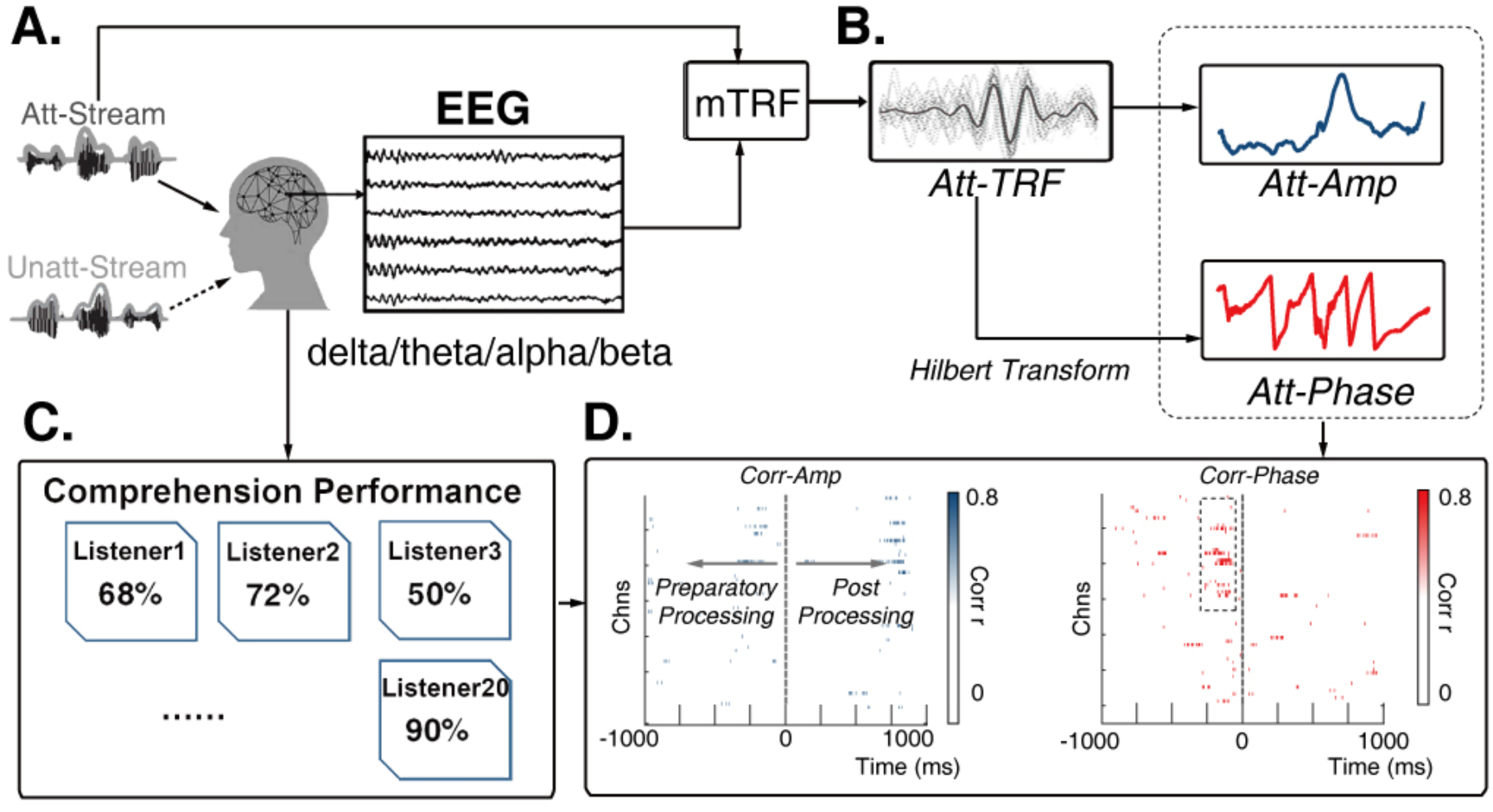
The analysis workflow. (A) The experimental paradigm. Participants attended to one of two simultaneously presented naturalistic, narrative speech streams while 60-channel EEG was recorded. (B) EEG data analysis. Neural responses were characterized using a TRF-based modeling method. The TRF-based neural responses were decomposed into the amplitude and the phase responses using the Hilbert transform. This procedure was conducted separately for attended (Att-) and unattended (Unatt-, not shown) speech streams, and separately for EEG data filtered at delta, theta, alpha and beta bands. (C) Comprehension performance. The participants completed a comprehension task after each speech comprehension trial. The average response accuracy over all trials per participant was taken as his/her comprehension performance. (D) Correlation analysis for comprehension performance-related neural responses. We calculated the correlation between either amplitude or phase responses and comprehension performance for each channel-latency bin. We defined neural activity before 0 ms as preparatory-processing and activity after 0 ms as post-processing. The results of the delta band were illustrated here. The colored channel-latency bins showed uncorrected significant correlation with comprehension performance. The dashed box in the ‘Corr-Phase’ plot indicates a significant channel-latency cluster by a cluster-based permutation test.

Prior to the modeling, the preprocessed EEG signals were re-referenced to the average of all scalp channels and then downsampled to 128 Hz. Then, the EEG data were filtered in delta (1–4 Hz), theta (4–8 Hz), alpha (8–12 Hz) and beta (12–30 Hz) bands (filter order: 64, one-pass forward filter). The use of a causal FIR filter ensured that filtered EEG signals were decided only by the current and previous data samples (de Cheveigné and Nelken 2019), which is important for the present research aim of preparatory speech processing. The filter order of 64 was chosen to keep a balance of temporal resolution and filter performance: the filtered EEG signals were therefore calculated based on the preceding 500 ms data (64 at 128 Hz).

The power envelopes of the speech signals were obtained using a Hilbert transform and then downsampled to the same sampling rate of 128 Hz. When denoting 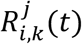 as the downsampled EEG signals from channel *i*, trial *k* filtered at one specific frequency band *j* (representing the four frequency bands) and *S*_*k*_(*t*) as the input speech power envelope corresponding to trial *k*, the corresponding neural response 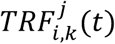 can be formulated as follows:

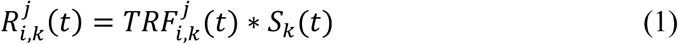

Where * represents the convolution operator. The latency in the neural response models 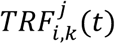 was set to vary from -1000 ms to 1000 ms post-stimulus. To control for overfitting, we varied the lambda from 10^−1^ to 10^3^ (lambda = 10^−1^, 10^0^, …, 10^3^) in the ridge regression (Di Liberto et al. 2015; Crosse et al. 2016; Broderick et al. 2018). The lambda value corresponding to the backward decoder that produced the highest cross-validated speech envelope reconstruction accuracy, averaged across trials, was selected as the regularization parameter for all trials per participant (Broderick et al. 2019). All TRF-based responses were further transformed into z-scores within the -1000 ms to 1000 ms time window for each channel separately per participant to account for across participants and across session differences (Pasley et al. 2012; Kleen et al. 2016). These z-scores were then used for the following analyses.

TRF models were calculated for attended and unattended speech processing separately using the corresponding speech streams as the input signal. TRF models were obtained for each EEG channel per participant, at the four frequency bands. It should be noted that we did not consider the input lateralization for the TRF models, as the observed behaviorally related findings were insensitive to the physical origin of the speech audios, but rather likely to reflect the lateralization of the human speech network. Fig. S1 provides the topographical information of our main results; it is similar to Fig. 2, but all results were calculated separately for speech stimuli from the left and right sides. The topographies were comparable.

**Figure 2.**
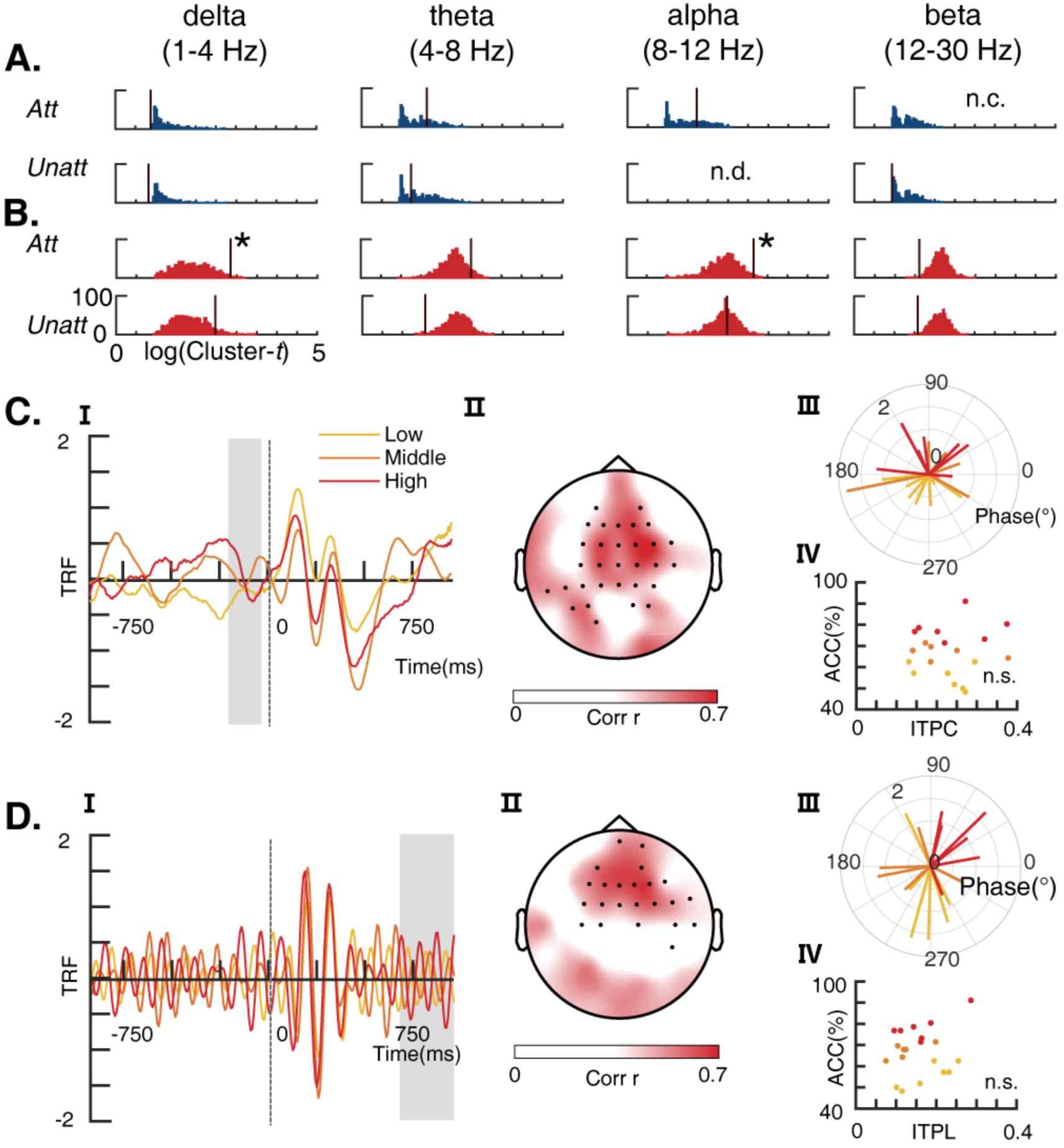
Speech comprehension performance related TRF-based responses to naturalistic speech. (A) Distributions of the cluster-level correlational t-statistics (log-transformation) from the 1, 000 permutated calculations of the correlation between the comprehension performance and the TRF-based amplitude response at the four frequency bands. The upper and lower panel for the attended and unattended responses, respectively. Vertical lines indicate the t-statistics of the clusters from the real data. N.C. means no cluster was formed and N.D. means the permutated distribution could not be generated (i.e., no cluster formed during the permutation calculation). (B) Distributions of the cluster-level correlational t-statistics from the 1, 000 permutated calculations of the correlation between the comprehension performance and the TRF-based phase response at the four frequency bands. The upper and lower panel for the attended and unattended responses, respectively. The two vertical lines with asterisks indicate statistically significant clusters from the real data. (C) Illustration of the significant phase-response cluster at the delta band. I. The time course of the TRFs at one representative channel (FC1). The three waveforms represent the average responses over the participants with comprehension performance of the attended speech ranking in the top (red), middle (orange), and bottom (yellow) tertiles (7, 6, and 7 participants, respectively). The shaded region depicts the time window of interest with a significant correlation. II. Topography of the average correlation *r*-values in the time window of interest. Black dots indicate the channels of interest in the cluster. III. The polar plot showing the average response at channel FC1 per participant. The vector lengths indicate the response amplitude and the vector directions indicate the phase angle. The three different colors show the comprehension performance rankings as in I. IV. The scatter plot showing the participants’ comprehension performance (accuracy in percentage) versus their inter-trial phase-locking (ITPL) values. The color of the dots indicates the comprehension performance rankings as in I and III. ‘n.s.’ means no significant correlation. (D) Illustration of the significant phase-response cluster at the alpha band. The plots in I and III are from channel FC1. The explanations of the sub-plots follow (C).

Amplitude and phase were calculated using the Hilbert transform of the TRF-based neural responses at the single-trial level. Hereby, the instantaneous amplitude 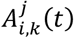 and the instantaneous phase 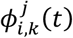 can be computed as

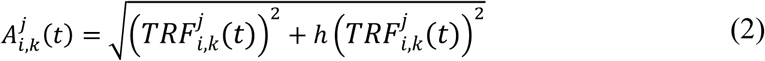

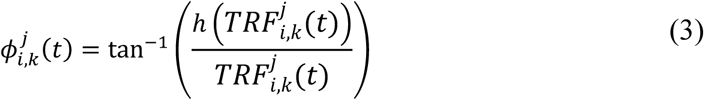

where 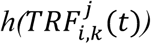 represents its Hilbert transform.

These single-trial instantaneous amplitude and phase were then averaged across all trials per participants to reflect one’s overall EEG responses to the naturalistic speech streams. Specifically, the single-trial amplitudes were averaged by taking their arithmetic mean value, whereas the single-trial phase responses were averaged by computing their circular mean (i.e., the mean phase angle).

In addition, the inter-trial phase locking (ITPL) was also calculated to evaluate the phase consistency across trials within each participant, as follows:

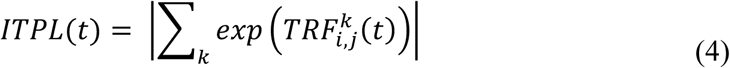

The phase-related ITPL value varies between 0 and 1; 0 refers to a situation in which the phase responses of different trials are uniformly distributed between 0 and 2π, and 1 means the phase responses from all trials are entirely locked to a fixed phase angle. The TRF analysis was conducted in MATLAB using the Multivariate Temporal Response Function (mTRF) toolbox (Crosse et al. 2016). All the other EEG processing procedures, as well as the statistical analyses, were conducted using the FieldTrip toolbox (Oostenveld et al. 2011).

### 2.7. Quantification and statistical analysis

The extracted TRF-based amplitude and phase responses were used to correlate with the speech comprehension performance of the attended speech at the participant level. The Spearman’s correlation was calculated between the amplitude response and the comprehension performance at each EEG channel and each individual latency across the participants. The correlations between the phase response and the comprehension performance were evaluated by computing the circular linear correlation using the CircStat toolbox (Berens 2009). Both the TRFs to the attended and unattended speech at the four frequency bands were included for this analysis.

Statistical analysis was performed to examine the significance of correlations over all channel-latency bins by computing the correlation *r*-values. To account for multiple comparisons, a nonparametric cluster-based permutation analysis was applied (Maris and Oostenveld 2007). In this procedure, neighboring channel-latency bins with an uncorrected correlational *p*-value below 0.01 were combined into clusters, for which the sum of the correlational *t*-statistics corresponding to the correlation *r*-values were obtained. A null-distribution was created through permutations of data across participants (*n* = 1,000 permutations), which defines the maximum cluster-level test statistics and corrected *p*-values for each cluster.

To investigate the attention modulation effect, we performed paired *t*-tests contrasting the TRFs to the attended speech versus the unattended speech. Both amplitude and phase were included in the analysis. The phase difference was calculated as the phase angle difference by the CircStat toolbox as well. A similar cluster-based permutation was used to control for the multiple comparison problem (*p* < .01 as the threshold, *n* = 1, 000 permutations).

### 2.8. Data and code availability statement

Dataset generated in our study has been uploaded to OSF (https://osf.io/87srv/?view_only=13f01e1f1f7b4cf98555ffacd878a53b). We have also provided data of individual participant, including the original EEG data and behavioral performance data.

Where existing toolboxes were used for data analysis, citations have been provided.

## 3. Results

### 3.1. Behavioral results

The average comprehension performance was significantly better for the 28 attended stories than for the 28 unattended stories (67.0±2.5% (standard error) vs. 36.0±1.6%, *t*(19) = 10.948, *p* < .001; the four-choice chance level: 25%). The participants reported a moderate level of attention (8.146±0.343 on a 10-point Likert scale) and attention difficulties (2.039±0.530 on a 10-point Likert scale). The accuracy for the attended story was significantly correlated with both the self-reported attention level (*r* = .476, *p* = .043) and attention difficulty (*r* = -.677, *p* = .001). The self-reported story familiarity level was low for all the participants (0.860±0.220 on a 10-point Likert scale) and was not correlated with comprehension performance (*r* = -.224, *p* = .342). These results suggest that participants’ selective attention was effectively manipulated, as well as good reliability of the measured comprehension performance. Most importantly, there was a large inter-individual difference in the participant-wise average comprehension performance for the attended stories; the response accuracy varied from 48.2% to 91.1%, which supports the feasibility of using these accuracy values as a behavioral indicator of comprehension-relevant neural signatures.

### 3.2. Speech comprehension performance related TRF-based responses

The nonparametric cluster-based permutation analysis revealed a significant correlation between the multi-channel amplitude and phase representation of the TRFs and individual speech comprehension performance of the attended speech. This corresponded to two clusters in the observed data (cluster-based permutation *p* < .05). Both the two significant clusters are reflected by TRF-based phase responses to attended speech, one at the delta band and the other at the alpha band (Figure 2A and 2B). No significant correlations were found for the TRFs to the unattended speech.

As shown in Figure 2C.I and II, the delta cluster is represented by TRF-based phase responses to attended speech at -200–0 ms relative to the onset of speech power envelope fluctuations over the fronto-central and left lateralized parietal regions (cluster-based permutation *p =* .012, mean circular linear correlation *r* = .787). The participants with their comprehension performance ranking in top, middle, and bottom tertiles were associated with different delta phase angles during this pre-speech-onset time window (average Φ_top_ = 73.411°, Φ_middle_ = 80.175°, and Φ_bottom_ = -134.767°, respectively, see Figure 2C.III). Specifically, the top-performing participants showed a negative peak within the time window of the cluster, whereas the bottom-performing participants showed a positive peak during this time. However, the participants’ ITPL did not significantly correlate with their comprehension performance (*r* = .127, *p* = . 593, Figure 2C.IV).

The alpha cluster was represented by TRF-based phase responses to attended speech at 650 – 980 ms post speech onset, with a fronto-central distribution (cluster-based permutation *p =* .031, mean *r* = .784, see Figure 2D.I and II). The individual difference in the phase angle of their responses was large, with average Φ_top_ = 41.904° for the top-performing participants and Φ_middle_ = -135.461°, Φ_bottom_ = -115.916° for the other two groups (Figure 2D.III). Again, there was no significant correlation between the ITPL and the comprehension performance (*r* = .267, *p* = .255, Figure 2D.IV).

### 3.3. Attention related TRF-based responses

The nonparametric cluster-based permutation analysis revealed a significant difference between the TRFs to the attended and unattended speech. The difference was manifested as a cluster involving a set of channels covering the left lateralized fronto-central and occipital regions (cluster-based permutation *p* < .001) at a latency of 240– 320 ms post the onset of speech power envelope fluctuations (Figure 3C). This attentional difference is reflected on TRF-based amplitude response at the theta band. No significant differences were observed in amplitude responses at other frequency bands and phase responses at all frequency bands (Figure 3A and 3B).

**Figure 3.**
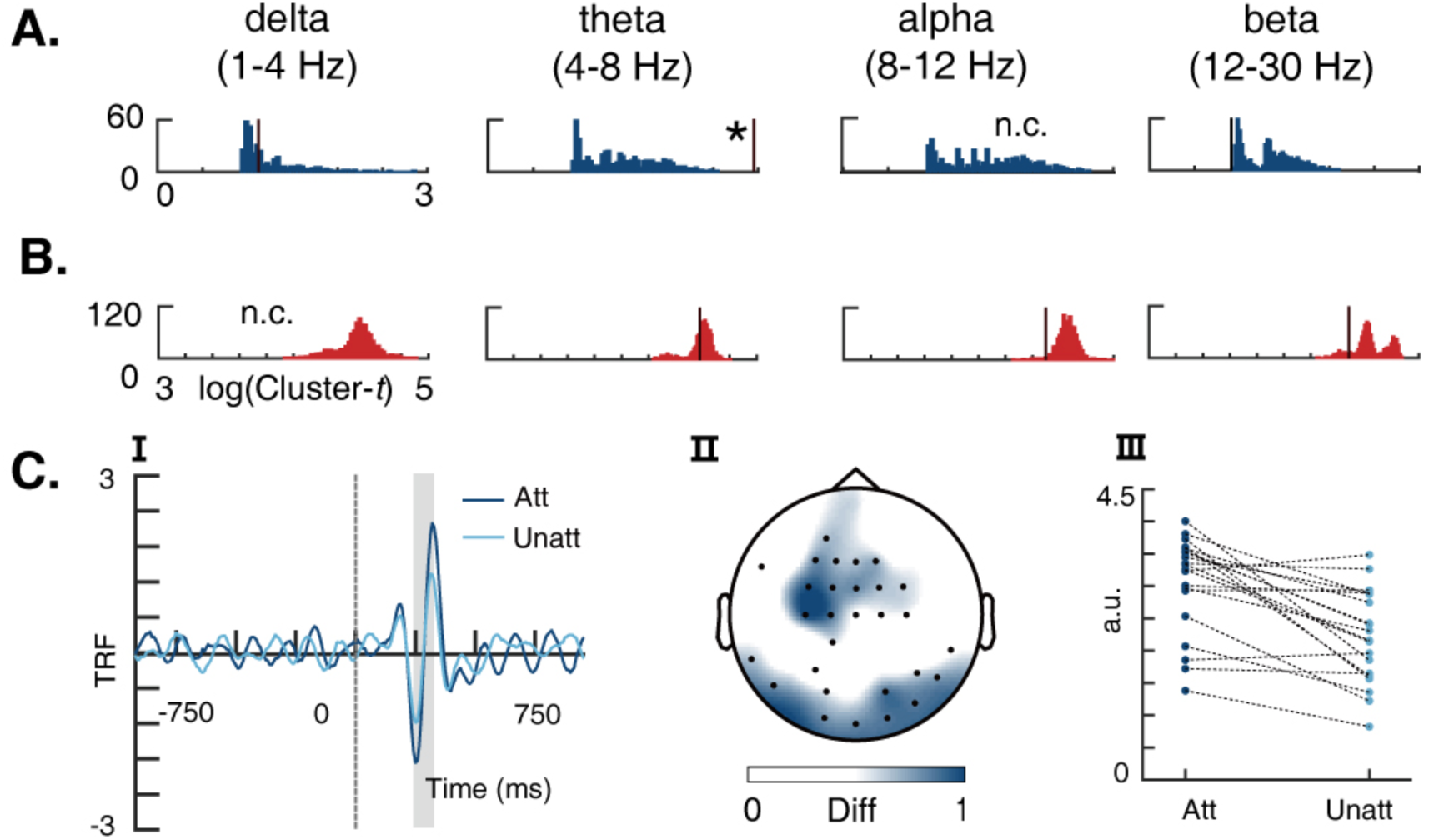
Attention related TRF-based responses to naturalistic speech. (A) Distributions of the cluster-level t-statistics (log-transformation) from the 1, 000 permutated calculations of the paired comparisons between the TRF-based attended and unattended amplitude responses at the four frequency bands. Vertical lines indicate the max cluster-level t-statistics in real data and the asterisk shows the statistically significant cluster from the real data. N.C. means no cluster was formed. (B) Distributions of the cluster-level t-statistics from the 1, 000 permutated calculations of the paired comparisons between the TRF-based attended and unattended phase responses at the four frequency bands. (C) Illustration of the significant amplitude-response cluster at the theta band. I. The time course of the TRFs at one representative channel (FC1). The two waveforms represent the average responses to the attended (dark blue) and unattended (light blue) speech. The shaded region depicts the time window of interest with significant differences between the two waveforms. II. Topography of the average amplitude response differences in the time window of interest. Black dots indicate the channels of interest in the cluster. III. The amplitude response difference per participant.

## 4. Discussion

The present study aimed to identify neural signatures that directly reflect the preparatory processing of upcoming speech. We used naturalistic narrative speech materials in a selective attention paradigm and a TRF-based approach for modeling the neural activity and observed preparatory neural activities before the onset of speech power envelope fluctuations. We found the phase responses to the attended speech at the delta band (1 – 4 Hz) were correlated with the comprehension performance of individual participants, with a latency of -200–0 ms before onset over the fronto-central and left lateralized parietal regions. The phase responses to the attended speech at the alpha band also correlated with comprehension performance, but with a latency of 650– 980 ms post onset over fronto-central regions. Distinct neural signatures were found for the attentional modulation, taking the form of TRF-based amplitude responses at a latency of 240–320 ms post onset over the left lateralized fronto-central and occipital regions. Our results provide direct neural evidence for how the brain prepares for the processing of upcoming speech.

Before detailed discussions, it is necessary to state that our assumption for a preparatory process is based on the observation that the TRF-based neural activities before the onset of speech power envelop fluctuations during the continuous speech stream were significantly correlated with comprehension performance. Recent TRF-based studies using naturalistic stimuli have reported reasonable latencies that resembled their ERP counterparts for describing selective auditory attention (∼200 ms) (Mirkovic et al. 2015; O’Sullivan et al. 2015), semantic violation processing (∼400 ms) (Broderick et al. 2018), and visual working memory (200–400 ms) (Huang et al. 2018). Although our findings have mainly focused on the window of < 0 ms, these studies support the rationale of using the TRF-based responses to reflect the time course of information processing in general. In addition, the present study also replicated the timing of previously reported attentional modulation of TRF-based responses to speech stimuli (Mirkovic et al. 2015; O’Sullivan et al. 2015). Therefore, the pre-onset latencies observed in the present study can be considered to represent a preparatory state that precedes speech processing. Moreover, the TRF method has enabled us to investigate the neural dynamics in an event-related-like manner but with naturalistic speech materials that are expected to better resemble real-world speech comprehension tasks.

Our results highlight in particular that the delta band phase at -200–0ms before the speech onset determines the comprehension accuracy of the listeners, which serves for the preparation of up comping speech information. As the analysis was performed on the neural responses to the to-be-processed speech at the pre-onset stage rather than those related to the preceding speech-related information (i.e., attentional cue, preceding words), our study has focused on a distinct processing phase other than previous studies that have mainly explored either the preparatory attention orientation and the speech-specific pre-activations by the preceding speech unit (DeLong et al. 2005; Söderström et al. 2016, 2018). Notably, the employment of the naturalistic speech materials has ensured sufficient variations of the interval between the preceding and the to-be-processed speech unit. Therefore, the present results are not likely to reflect the neural responses to the preceding speech unit. Although there was one study that has reported a sustained difference in the neural activities at around 400 ms prior to the onset of the upcoming speech (Lee et al. 2013), their ‘pre-onset’ activities mainly reflected the preparatory attention orientation by a visually-presented abstract attentional cue. In contrast, the results of the present study are expected to reflect how the human brain makes use of the rich contextual information in the naturalistic speech materials to infer and prepare for the upcoming speech information.

The timing and the spatial distribution of the reported pre-onset preparatory activity is better compared with the studies on pre-stimulus oscillatory activities. Visual pre-stimulus studies have generally reported perceptual-relevant neural activities with timings of 100–400 ms prior to stimulus onset (Harris et al. 2018; Wöstmann et al. 2019). In our research, we have found a similar time window for those studies. Our results, therefore, suggest that approximately -200–0 ms prior to stimulus onset might have a general implication for the preparation of sensory information processing. As the present study employed naturalistic (speech) materials that are expected to provide much richer contextual information as compared to the simple and abstract stimuli in most of the previous studies, our results might provide more reliable support for the observed timing on preparatory sensory processing. The result is also in line with a recent study, which indicated that the delta band entrainment at 100 ms before the stimulus is related to the noise-induced comprehension difference (Etard and Reichenbach 2019). Meanwhile, the central-parietal regions found in our study involved in the preparatory processing are also consistent with the previous auditory pre-stimulus studies (Stefanics et al. 2010; Kayser et al. 2016). The central-parietal responses could be related to the predictive processing of speech meaning and could recruit a mechanism that is similar to that underlying the classical central-parietal N400 response (Federmeier 2007; Lau et al. 2008; Szewczyk and Schriefers 2018) or possibly preparatory attentional orientation as well (Holmes et al. 2016). The left lateralized parietal regions could also be linked to the speech-specific processing, e.g. the Wernicke’s area (Hickok and Poeppel 2007).

Our study expands on findings from the studies about the functional role of the delta band in speech preparatory processing. The delta band could help the segmentation or identification of intonation phrases, which is essential for the preparation and prediction of upcoming speech (Giraud and Poeppel 2012; Ding et al. 2015; Kösem et al. 2018; Meyer 2018; Morillon et al. 2019). In particular, the phase of the delta band before the stimulus is related to the hit rate afterward (Ng et al. 2012), or the behavioral consequence (i.e., the reaction time) in the auditory studies using simple, isolated stimulus (Lakatos et al. 2008; Stefanics et al. 2010; Henry and Obleser 2012) . It is in contrast to visual studies in which the alpha band is mostly pronounced in the pre-stimulus stage (Busch et al. 2009; Mathewson et al. 2011; Milton and Pleydell-Pearce 2016), suggesting active auditory perception is dominated by lower-frequency (Ng et al. 2012; VanRullen 2016). In addition, the brain activity dynamically tracks speech streams using the delta phase (Zion Golumbic, Cogan, et al. 2013) and temporal predictions are encoded by delta neural oscillations (Morillon and Baillet 2017; Auksztulewicz et al. 2018). Our findings are in line with the previous studies about the delta band phase’s role in preparatory processing and unambiguously show that the comprehension performance could be elevated when the delta is better prepared at a particular phase angle.

The neural mechanisms of the preparatory process were investigated using correlation analysis of the TRF-based neural activity. In line with recent TRF-based studies, we observed attention-related neural responses (Mirkovic et al. 2015, 2016; O’Sullivan et al. 2015), with the peak attention effect represented by theta amplitude activities at 250– 320 ms post-stimulus onset over the central and occipital regions. In contrast, the comprehension-related post-onset neural signatures were in the alpha band at 670–980 ms. The result could be interpreted for a functional role of alpha-band for a top-down control mechanism to achieve the preparatory process, as the phase of alpha oscillations has an active role in attentional temporal predictions (Händel et al. 2011; Bonnefond and Jensen 2012; Samaha et al. 2015). Meanwhile, in auditory studies, the alpha frequency band has also been suggested to be associated with working memory (Bonnefond and Jensen 2012; Meyer 2018), capable of storing semantics in sentences (Haarmann and Cameron 2005) and syntax information (Bonhage et al. 2017) and lexical decision (Strauß et al. 2015), which are also closely related to speech comprehension.

The neural mechanisms of the preparatory process were further explored by inspecting their relationship with attention. While our results are in line with previous research that has reported low-frequency phase track the envelope of attended speech (Ding and Simon 2012; Zion Golumbic, Ding, et al. 2013), we provide further evidence on how such interactions could affect behavior (i.e., comprehension). Indeed, we only found the correlation between the attended TRF-based neural activities and the comprehension performance in the pre-onset stage, suggesting a possible attentional facilitation of preparation of speech processing.

This study has some limitations that should be noted. While the use of naturalistic speech materials could better resemble the real-world speech comprehension scenarios, it is difficult to infer which specific types of information (e.g. timing, phoneme, etc.) were the main contributor for the observed preparatory activities. The distributed brain regions involved in the preparatory process may provide a guidance for designing further experiments to have an in-depth exploration. The present study used the speech power envelope as the reference signal from which the TRF models were derived, which could reflect the speech information at all linguistic levels due to the highly redundant information shared across levels (Di Liberto et al. 2015; Daube et al. 2019). While such an operation has the advantage of providing a general overview about preparatory processing, further investigations are necessary to differentiate possible contributions at different linguistic levels (Di Liberto et al. 2015; Broderick et al. 2018). Meanwhile, caution must be taken when interpreting the timing of the preparatory activities. While the preparatory activity as early as ∼200 ms before speech onset could be the result of an optimized utilization of the rich contextual information provided by the naturalistic speech materials, such timings may be dependent upon the materials per se. Further studies are necessary to investigate the possible material dependence of these timings, for instance, by employed an extended amount of speech materials. Also, denser MEG recordings together with source localization methods are expected to more precisely identify the brain regions for preparatory speech processing (Mazaheri et al. 2009; Nolte and Müller 2010). Besides, an inter-individual level regression analysis method was chosen, as the average comprehension questionnaire accuracies across all stories within each participant were believed to provide a more reliable estimation of the speech comprehension performance than the single-trial accuracies. Thus, our results do not necessarily imply that the observed neural signatures reflect the participants’ trait-like, stable speech processing style. Alternatively, it could be more plausible to consider these neural signatures to reflect a more or less efficient speech processing state. More theoretical and empirical research is needed to clarify the underlying mechanisms.

## Supporting information

**Figure S1.**
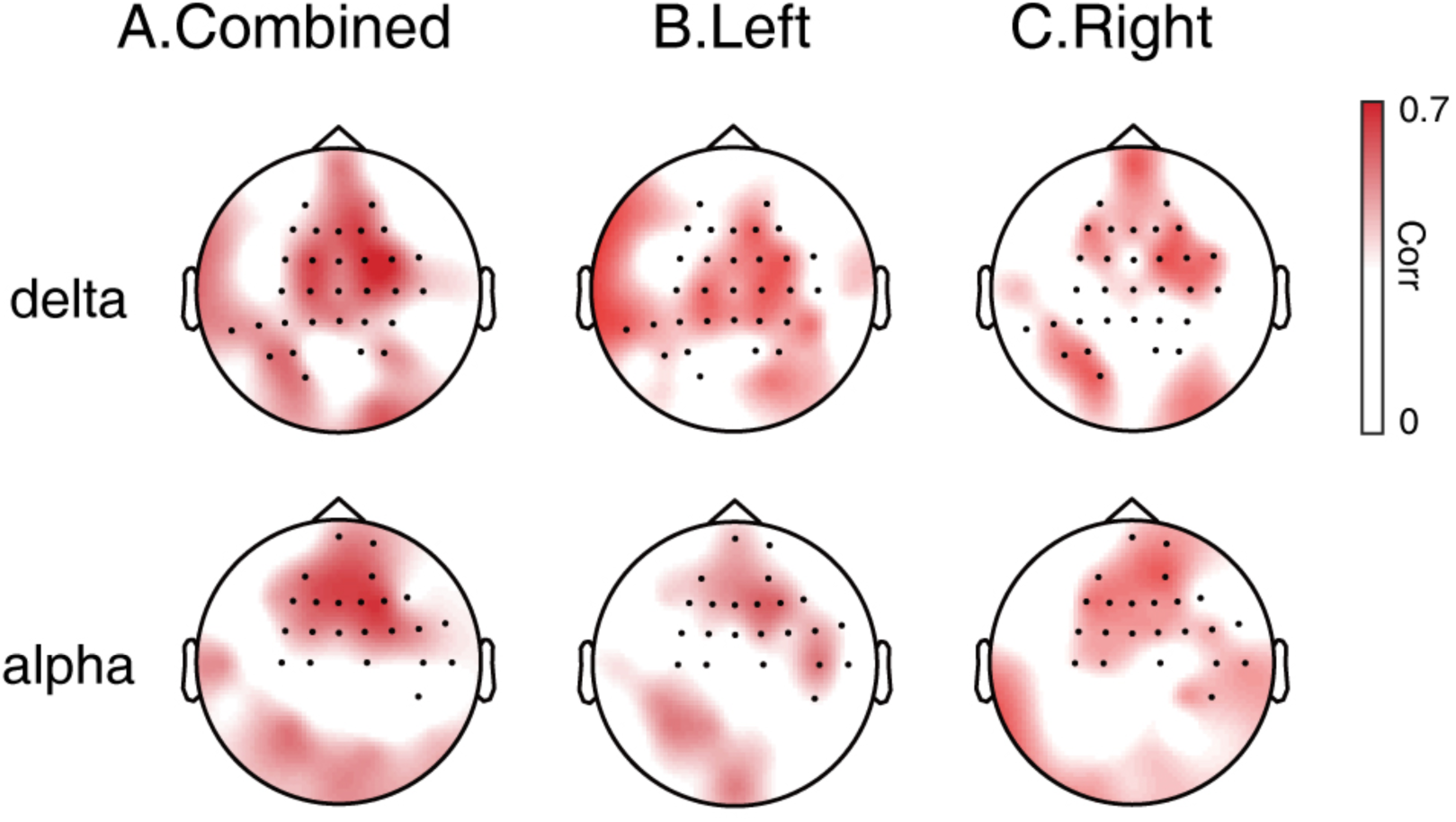
Topographies of the two responses as shown in Figure 2 (A) and calculated separately for the speech stimuli delivered to the left side (B) and the right side (C) only.

## Acknowledgments

This work was supported by the National Natural Science Foundation of China (NSFC) and the German Research Foundation (DFG) in project Crossmodal Learning (grant number: NSFC 61621136008/DFG TRR-169/C1, B1), the National Key Research and Development Plan (grant number: 2016YFB1001200), the National Natural Science Foundation of China (grant number: 61977041 and U1736220), the National Social Science Foundation of China (grant number: 17ZDA323) and Tsinghua University Initiative Scientific Research Program (grant number: 20197010006).

The authors would like to thank Prof. Dr. Xiaoqin Wang and Dr. Yue Ding for providing the shielded room for the experiment as well as necessary technical support.

## References

Alexandrou AM, Saarinen T, Kujala J, Salmelin R. 2020. Cortical entrainment: what we can learn from studying naturalistic speech perception. Lang Cogn Neurosci. 35:681–693.

Arnal LH, Wyart V, Giraud AL. 2011. Transitions in neural oscillations reflect prediction errors generated in audiovisual speech. Nat Neurosci. 14:797–801.

Auksztulewicz R, Friston K. 2015. Attentional enhancement of auditory mismatch responses: A DCM/MEG study. Cereb Cortex. 25:4273–4283.

Auksztulewicz R, Schwiedrzik CM, Thesen T, Doyle W, Devinsky O, Nobre AC, Schroeder CE, Friston KJ, Melloni L. 2018. Not all predictions are equal: “what” and “when” predictions modulate activity in auditory cortex through different mechanisms. J Neurosci. 38:8680–8693.

Avramiea AE, Hardstone R, Lueckmann JM, Bim J, Mansvelder HD, Linkenkaer-Hansen K. 2020. Pre-stimulus phase and amplitude regulation of phase-locked responses is maximized in the critical state. Elife. 9:1–17.

Bednar A, Lalor EC. 2018. Neural tracking of auditory motion is reflected by delta phase and alpha power of EEG. Neuroimage. 181:683–691.

Bednar A, Lalor EC. 2020. Where is the cocktail party? Decoding locations of attended and unattended moving sound sources using EEG. Neuroimage. 205:116283.

Berens P. 2009. CircStat : A MATLAB Toolbox for Circular Statistics. J Stat Softw. 31:293–295.

Bonhage CE, Meyer L, Gruber T, Friederici AD, Mueller JL. 2017. Oscillatory EEG dynamics underlying automatic chunking during sentence processing. Neuroimage. 152:647–657.

Bonnefond M, Jensen O. 2012. Alpha oscillations serve to protect working memory maintenance against anticipated distracters. Curr Biol. 22:1969–1974.

Brainard DH, Brainard DH. 1997. The Psychophysics Toolbox. In: Spatial vision. p. 433–436.

Broderick MP, Anderson AJ, Di Liberto GM, Crosse MJ, Lalor EC. 2018. Electrophysiological Correlates of Semantic Dissimilarity Reflect the Comprehension of Natural, Narrative Speech. Curr Biol. 28:803-809.e3.

Broderick MP, Anderson AJ, Lalor EC. 2019. Semantic Context Enhances the Early Auditory Encoding of Natural Speech. J Neurosci. 39:7564–7575.

Busch NA, Dubois J, VanRullen R. 2009. The phase of ongoing EEG oscillations predicts visual perception. J Neurosci. 29:7869–7876.

Cherry EC. 1953. Some experiments on the recognition of speech, with one and with two ears. J Acoust Soc Am. 25:975–979.

Crosse MJ, Di Liberto GM, Bednar A, Lalor EC. 2016. The Multivariate Temporal Response Function (mTRF) Toolbox: A MATLAB Toolbox for Relating Neural Signals to Continuous Stimuli. Front Hum Neurosci. 10:604.

Daube C, Ince RAA, Gross J. 2019. Simple Acoustic Features Can Explain Phoneme-Based Predictions of Cortical Responses to Speech. Curr Biol. 29:1924-1937.e9.

de Cheveigné A, Nelken I. 2019. Filters: When, Why, and How (Not) to Use Them. Neuron. 102: 280–293.

DeLong KA, Urbach TP, Kutas M. 2005. Probabilistic word pre-activation during language comprehension inferred from electrical brain activity. Nat Neurosci. 8:1117–1121.

DeWitt I, Rauschecker JP. 2012. Phoneme and word recognition in the auditory ventral stream. Proc Natl Acad Sci U S A. 109:505–514.

Di Liberto GM, O’Sullivan JA, Lalor EC. 2015. Low-frequency cortical entrainment to speech reflects phoneme-level processing. Curr Biol. 25:2457–2465.

Dikker S, Pylkkänen L. 2013. Predicting language: MEG evidence for lexical preactivation. Brain Lang. 127:55–64.

Ding N, Melloni L, Zhang H, Tian X, Poeppel D. 2015. Cortical tracking of hierarchical linguistic structures in connected speech. Nat Neurosci. 19:158–164.

Ding N, Simon JZ. 2012. Emergence of neural encoding of auditory objects while listening to competing speakers. Proc Natl Acad Sci U S A. 109:11854–11859.

Donhauser PW, Baillet S. 2020. Two Distinct Neural Timescales for Predictive Speech Processing. Neuron. 105:385-393.e9.

ElShafei HA, Bouet R, Bertrand O, Bidet-Caulet A. 2018. Two Sides of the Same Coin: Distinct Sub-Bands in the α Rhythm Reflect Facilitation and Suppression Mechanisms during Auditory Anticipatory Attention. eneuro. 5:1–14.

Engel AK, Gerloff C, Hilgetag CC, Nolte G. 2013. Intrinsic Coupling Modes: Multiscale Interactions in Ongoing Brain Activity. Neuron. 80:867–886.

Etard O, Reichenbach T. 2019. Neural Speech Tracking in the Theta and in the Delta Frequency Band Differentially Encode Clarity and Comprehension of Speech in Noise. J Neurosci. 39:5750–5759.

Federmeier KD. 2007. Thinking ahead: The role and roots of prediction in language comprehension. Psychophysiology. 44:491–505.

Friederici AD. 2012. The cortical language circuit: From auditory perception to sentence comprehension. Trends Cogn Sci. 16:262–268.

Fries P. 2015. Rhythms for Cognition: Communication through Coherence. Neuron. 88:220–235.

Friston K. 2005. A theory of cortical responses. Philos Trans R Soc B Biol Sci. 360:815–836.

Galindo-Leon EE, Stitt I, Pieper F, Stieglitz T, Engler G, Engel AK. 2019. Context-specific modulation of intrinsic coupling modes shapes multisensory processing. Sci Adv. 5:1–13.

Giraud AL, Poeppel D. 2012. Cortical oscillations and speech processing: Emerging computational principles and operations. Nat Neurosci. 15:511–517.

Haarmann HJ, Cameron KA. 2005. Active maintenance of sentence meaning in working memory : Evidence from EEG coherences. 57:115–128.

Händel BF, Haarmeier T, Jensen O. 2011. Alpha oscillations correlate with the successful inhibition of unattended stimuli. J Cogn Neurosci. 23:2494–2502.

Harris AM, Dux PE, Mattingley JB. 2018. Detecting Unattended Stimuli Depends on the Phase of Prestimulus Neural Oscillations. J Neurosci. 38:3092–3101.

Henry MJ, Obleser J. 2012. Frequency modulation entrains slow neural oscillations and optimizes human listening behavior. Proc Natl Acad Sci. 109:20095–20100.

Hickok G. 2012. Computational neuroanatomy of speech production. Nat Rev Neurosci. 13:135–145.

Hickok G, Houde J, Rong F. 2011. Sensorimotor Integration in Speech Processing: Computational Basis and Neural Organization. Neuron. 69:407–422.

Hickok G, Poeppel D. 2007. The cortical organization of speech processing. Nat Rev Neurosci. 8:393–402.

Hill KT, Miller LM. 2010. Auditory attentional control and selection during cocktail party listening. Cereb Cortex. 20:583–590.

Hisagi M, Shafer VL, Strange W, Sussman ES. 2015. Neural measures of a Japanese consonant length discrimination by Japanese and American English listeners: Effects of attention. Brain Res. 1626:218–231.

Holmes E, Kitterick PT, Summerfield AQ. 2016. EEG activity evoked in preparation for multi-talker listening by adults and children. Hear Res. 336:83–100.

Holmes E, Kitterick PT, Summerfield AQ. 2018. Cueing listeners to attend to a target talker progressively improves word report as the duration of the cue-target interval lengthens to 2,000 ms. Attention, Perception, Psychophys. 80:1520–1538.

Huang Q, Jia J, Han Q, Luo H. 2018. Fast-backward replay of sequentially memorized items in humans. Elife. 7:e35164.

Iemi L, Busch NA, Laudini A, Haegens S, Samaha J, Villringer A, Nikulin V V. 2019. Multiple mechanisms link prestimulus neural oscillations to sensory responses. Elife. 8:e43620.

Jehee JFM, Ballard DH. 2009. Predictive Feedback Can Account for Biphasic Responses in the Lateral Geniculate Nucleus. PLoS Comput Biol. 5:e1000373.

Kayser SJ, McNair SW, Kayser C. 2016. Prestimulus influences on auditory perception from sensory representations and decision processes. Proc Natl Acad Sci. 113:4842–4847.

Kleen JK, Testorf ME, Roberts DW, Scott RC, Jobst BJ, Holmes GL, Lenck-Santini PP. 2016. Oscillation phase locking and late ERP components of intracranial hippocampal recordings correlate to patient performance in a working memory task. Front Hum Neurosci. 10:1–14.

Kok P, Jehee JFM, de Lange FP. 2012. Less Is More: Expectation Sharpens Representations in the Primary Visual Cortex. Neuron. 75:265–270.

Kok P, Mostert P, De Lange FP. 2017. Prior expectations induce prestimulus sensory templates. Proc Natl Acad Sci U S A. 114:10473–10478.

Kösem A, Bosker HR, Takashima A, Meyer A, Jensen O, Hagoort P. 2018. Neural Entrainment Determines the Words We Hear. Curr Biol. 28:2867-2875.e3.

Kutas M, Federmeier KD. 2011. Thirty Years and Counting: Finding Meaning in the N400 Component of the Event-Related Brain Potential (ERP). Annu Rev Psychol. 62:621–647.

Lakatos P, Karmos G, Mehta AD, Ulbert I, Schroeder CE. 2008. Entrainment of Neuronal Oscillations as a Mechanism of Attentional Selection. Science. 320:110–113.

Lalor EC, Pearlmutter BA, Reilly RB, McDarby G, Foxe JJ. 2006. The VESPA: A method for the rapid estimation of a visual evoked potential. Neuroimage. 32:1549–1561.

Lau EF, Phillips C, Poeppel D. 2008. A cortical network for semantics: (de)constructing the N400. Nat Rev Neurosci. 9:920–933.

Lee AKC, Rajaram S, Xia J, Bharadwaj H, Larson E, Hämäläinen MS, Shinn-Cunningham BG. 2013. Auditory selective attention reveals preparatory activity in different cortical regions for selection based on source location and source pitch. Front Neurosci. 6:1–9.

Maris E, Oostenveld R. 2007. Nonparametric statistical testing of EEG- and MEG-data. J Neurosci Methods. 164:177–190.

Marzecová A, Widmann A, SanMiguel I, Kotz SA, Schröger E. 2017. Interrelation of attention and prediction in visual processing: Effects of task-relevance and stimulus probability. Biol Psychol. 125:76–90.

Mathewson KE, Lleras A, Beck DM, Fabiani M, Ro T, Gratton G. 2011. Pulsed out of awareness: EEG alpha oscillations represent a pulsed-inhibition of ongoing cortical processing. Front Psychol. 2:1–15.

Mazaheri A, Nieuwenhuis ILC, Van Dijk H, Jensen O. 2009. Prestimulus alpha and mu activity predicts failure to inhibit motor responses. Hum Brain Mapp. 30:1791–1800.

Meyer L. 2018. The neural oscillations of speech processing and language comprehension: state of the art and emerging mechanisms. Eur J Neurosci. 48:2609–2621.

Milton A, Pleydell-Pearce CW. 2016. The phase of pre-stimulus alpha oscillations influences the visual perception of stimulus timing. Neuroimage. 133:53–61.

Mirkovic B, Bleichner MG, De Vos M, Debener S. 2016. Target speaker detection with concealed EEG around the ear. Front Neurosci. 10:1–11.

Mirkovic B, Debener S, Jaeger M, Vos M De. 2015. Decoding the attended speech stream with multi-channel EEG: implications for online, daily-life applications. J Neural Eng. 12:046007.

Morillon B, Arnal LH, Schroeder CE, Keitel A. 2019. Prominence of delta oscillatory rhythms in the motor cortex and their relevance for auditory and speech perception. Neurosci Biobehav Rev. 107:136–142.

Morillon B, Baillet S. 2017. Motor origin of temporal predictions in auditory attention. Proc Natl Acad Sci. 114:E8913–E8921.

Ng BSW, Schroeder T, Kayser C. 2012. A Precluding But Not Ensuring Role of Entrained Low-Frequency Oscillations for Auditory Perception. J Neurosci. 32:12268–12276.

Nolden S, Ibrahim CN, Koch I. 2019. Cognitive control in the cocktail party: Preparing selective attention to dichotically presented voices supports distractor suppression. Attention, Perception, Psychophys. 81:727–737.

Nolte G, Müller KR. 2010. Localizing and estimating causal relations of interacting brain rhythms. Front Hum Neurosci. 4:1–5.

O’Sullivan JA, Power AJ, Mesgarani N, Rajaram S, Foxe JJ, Shinn-Cunningham BG, Slaney M, Shamma SA, Lalor EC. 2015. Attentional Selection in a Cocktail Party Environment Can Be Decoded from Single-Trial EEG. Cereb Cortex. 25:1697–1706.

Oostenveld R, Fries P, Maris E, Schoffelen J. 2011. FieldTrip: Open Source Software for Advanced Analysis of MEG, EEG, and Invasive Electrophysiological Data. Comput Intell Neurosci. 2011:1–9.

Pasley BN, David S V., Mesgarani N, Flinker A, Shamma SA, Crone NE, Knight RT, Chang EF. 2012. Reconstructing Speech from Human Auditory Cortex. PLoS Biol. 10:e1001251.

Pisoni DB, Luce PA. 1987. Acoustic-phonetic representations in word recognition. Cognition. 25:21–52.

Rao RPN, Ballard DH. 1999. Predictive coding in the visual cortex: A functional interpretation of some extra-classical receptive-field effects. Nat Neurosci. 2:79–87.

Rassi E, Wutz A, Müller-Voggel N, Weisz N. 2019. Prestimulus feedback connectivity biases the content of visual experiences. Proc Natl Acad Sci. 116:16056–16061.

Samaha J, Bauer P, Cimaroli S, Postle BR. 2015. Top-down control of the phase of alpha-band oscillations as a mechanism for temporal prediction. Proc Natl Acad Sci. 112:8439–8444.

Schröger E, Kotz SA, SanMiguel I. 2015. Bridging prediction and attention in current research on perception and action. Brain Res. 1626:1–13.

Schröger E, Marzecová A, Sanmiguel I. 2015. Attention and prediction in human audition: A lesson from cognitive psychophysiology. Eur J Neurosci. 41:641–664.

Smith JL, Johnstone SJ, Barry RJ. 2006. Effects of pre-stimulus processing on subsequent events in a warned Go/NoGo paradigm: Response preparation, execution and inhibition. Int J Psychophysiol. 61:121–133.

Smout CA, Tang MF, Garrido MI, Mattingley JB. 2019. Attention promotes the neural encoding of prediction errors. PLoS Biol. 17:1–22.

Söderström P, Horne M, Frid J, Roll M. 2016. Pre-Activation Negativity (PrAN) in Brain Potentials to Unfolding Words. Front Hum Neurosci. 10:1–11.

Söderström P, Horne M, Mannfolk P, van Westen D, Roll M. 2018. Rapid syntactic pre-activation in Broca’s area: Concurrent electrophysiological and haemodynamic recordings. Brain Res. 1697:76–82.

Sonkusare S, Breakspear M, Guo C. 2019. Naturalistic Stimuli in Neuroscience: Critically Acclaimed. Trends Cogn Sci. 1–16.

Stefanics G, Hangya B, Hernadi I, Winkler I, Lakatos P, Ulbert I. 2010. Phase Entrainment of Human Delta Oscillations Can Mediate the Effects of Expectation on Reaction Speed. J Neurosci. 30:13578–13585.

Strauß XA, Henry MJ, Scharinger XM, Obleser XJ. 2015. Alpha Phase Determines Successful Lexical Decision in Noise. 35:3256–3262.

Szewczyk JM, Schriefers H. 2018. The N400 as an index of lexical preactivation and its implications for prediction in language comprehension. Lang Cogn Neurosci. 33:665–686.

Teng X, Ma M, Yang J. 2020. Constrained Structure of Ancient Chinese Poetry Facilitates Speech Content Grouping. Curr Biol. 30:1–7.

Tian X, Ding N, Teng X, Bai F, Poeppel D. 2018. Imagined speech influences perceived loudness of sound. Nat Hum Behav. 2:225–234.

Van Dijk H, Schoffelen JM, Oostenveld R, Jensen O. 2008. Prestimulus oscillatory activity in the alpha band predicts visual discrimination ability. J Neurosci. 28:1816–1823.

Van Petten C, Luka BJ. 2012. Prediction during language comprehension: Benefits, costs, and ERP components. Int J Psychophysiol. 83:176–190.

VanRullen R. 2016. Perceptual Cycles. Trends Cogn Sci. 20:723–735.

Verhulst S, Altoè A, Vasilkov V. 2018. Computational modeling of the human auditory periphery: Auditory-nerve responses, evoked potentials and hearing loss. Hear Res. 360:55–75.

Wang L, Kuperberg G, Jensen O. 2018. Specific lexico-semantic predictions are associated with unique spatial and temporal patterns of neural activity. Elife. 7:1–24.

Wöstmann M, Waschke L, Obleser J. 2019. Prestimulus neural alpha power predicts confidence in discriminating identical auditory stimuli. Eur J Neurosci. 49:94–105.

Zion Golumbic E, Cogan GB, Schroeder CE, Poeppel D. 2013. Visual input enhances selective speech envelope tracking in auditory cortex at a “cocktail party”. J Neurosci. 33:1417–1426.

Zion Golumbic EM, Ding N, Bickel S, Lakatos P, Schevon CA, McKhann GM, Goodman RR, Emerson R, Mehta AD, Simon JZ, Poeppel D, Schroeder CE. 2013. Mechanisms underlying selective neuronal tracking of attended speech at a “cocktail party.” Neuron. 77:980–991.

Zoefel B, Archer-Boyd A, Davis MH. 2018. Phase Entrainment of Brain Oscillations Causally Modulates Neural Responses to Intelligible Speech. Curr Biol. 28:401-408.e5.

